# Chromosome-scale genome assembly of a Japanese chili pepper landrace, *Capsicum annuum* ‘Takanotsume’

**DOI:** 10.1101/2022.09.30.510245

**Authors:** Kenta Shirasawa, Munetaka Hosokawa, Yasuo Yasui, Atsushi Toyoda, Sachiko Isobe

**Affiliations:** Department of Frontier Research and Development, Kazusa DNA Research Institute, Kisarazu, Japan; Department of Agriculture, Kindai University, Nara, Japan; Agricultural Technology and Innovation Research Institute, Kindai University, Nara, Japan; Graduate School of Agriculture, Kyoto University, Kyoto, Japan; Advanced Genomics Center, National Institute of Genetics, Mishima, Japan

## Abstract

Here, we report the genome sequence of a popular Japanese chili pepper landrace, *Capsicum annuum* ‘Takanotsume’. We used long-read sequencing and optical mapping, together with the genetic mapping technique, to obtain the chromosome-scale genome assembly of ‘Takanotsume’. The assembly consists of 12 pseudomolecules, which corresponds to the basic chromosome number of *C. annuum*, and is 3,058.5 Mb in size, spanning 97.0% of the estimated genome size. A total of 34,324 high-confidence genes were predicted in the genome, and 83.4% of the genome assembly was occupied by repetitive sequences. Comparative genomics of linked-read sequencing-derived *de novo* genome assemblies of two *Capsicum chinense* lines and whole-genome resequencing-derived genome assemblies of *Capsicum* species revealed not only nucleotide sequence variations but also genome structure variations (i.e., chromosomal rearrangements) between ‘Takanotsume’ and its relatives. Overall, the genome sequence data generated in this study will accelerate the pan-genomics and breeding of *Capsicum*, and facilitate the dissection of genetic mechanisms underlying the agronomically important traits of ‘Takanotsume’.

## Introduction

The genus *Capsicum* includes four major species, *C. annuum, C. baccatum, C. chinense*, and *C. frutescens*, all of which are used as vegetables and spices^1^. Because of partial cross-compatibility among *Capsicum* species, attractive cultivars have been bred worldwide through both inter- and intraspecific crossing^1^. Therefore, pedigrees of interspecific hybrids are complicated and error-prone during the breeding process. The availability of interspecific hybrids depends on the combinations of parental lines used for their generation^2^. Some combinations generate morphologically abnormal F_1_ hybrids, which fail to survive^3^. This phenomenon is caused by a negative interaction between two independent genetic loci, a hypothesis also known as the Bateson–Dobzhansky–Muller (BDM) model, which has been observed in wide interspecific crosses in animals and plants, including pepper^4^.

To the best of our knowledge, the genomes of four *C. annuum* lines, one *C. baccatum* line, and one *C. chinense* line have been sequenced to date^5–10^. These sequences were constructed using two next-generation sequencing technologies: short-read sequencing and the error-prone long-read sequencing. Since the genomes of *Capsicum* species are larger and more complex than those of their relatives, e.g., *Solanum* species^11–13^, complete and high-quality genome sequencing of *Capsicum* might be difficult with the existent technologies. Therefore, the available sequence data have gaps, even though the sequences are assembled at the chromosome level^5–10^. Recent advances in sequencing technologies enable the generation of high-quality long reads, also known as HiFi reads^14^. Furthermore, techniques such as chromosome conformation capture^15^, which generates chromatin contact maps, and optical mapping^16^, which outputs high-resolution genome-wide restriction maps, are also available. These technologies could be used to assemble the genomes of multiple lines of different *Capsicum* species, generating the *Capsicum* pan-genome^17^, which would enhance our understanding of its genetic mechanisms and provide insights into *Capsicum* evolution.

‘Takanotsume’ (which in Japanese literally translates to “The Claw of the Hawk”) is a Japanese *C. annuum* landrace named after the shape of its fruit, which is similar to that of the nails of hawks. ‘Takanotsume’ plants exhibit indeterminate growth, with spread-out branching habit, and are cultivated for the thin-fleshed fruits^18^, which are used as a spice with a pungency level of approximately 11,900 on the Scoville scale^19^. Because of the rapid water loss from its fruits post-harvest, ‘Takanotsume’ has become a popular cultivar for spice purposes, and its derivative lines, such as ‘Hontaka’ and ‘Daruma’, have been distributed all over Japan^18^. However, the pedigree of ‘Takanotsume’ is unclear. ‘Takanotsume’ also possesses some unique characteristics, including two independent genes, which confer interspecific cross-compatibility explained by the BDM model^3^, and high ribonuclease activity in leaves, which could combat chrysanthemum stunt viroid *in vivo*^20^.

To reveal the genetic mechanisms underlying the attractive traits of ‘Takanotsume’, a high-quality genome assembly is required. In this study, we employed the HiFi sequencing technology, together with optical mapping and genetic mapping methods, to generate a chromosome-scale genome sequence assembly of ‘Takanotsume’. Comparative genomics revealed nucleotide sequence and chromosome structural variations within the *Capsicum* species. The genome sequence and variant information obtained in this study would be helpful for elucidating the genetic mechanisms controlling the unique traits of ‘Takanotsume’.

## Materials and methods

### Plant materials

*Capsicum annuum* landrace ‘Takanotsume’, which is maintained through self-pollination at Department of Agriculture, Kindai University, Nara, Japan, as well as thirteen *Capsicum* lines, including six *C. annuum* lines (‘106’, ‘110’, ‘Sweet Banana’, ‘California Wonder’, ‘Murasaki’, and ‘Nikko’), two *C. baccatum* lines (‘28’ and ‘Aji Rojo’), and five *C. chinense* lines (‘3686’, ‘3687’, ‘Charapita’, ‘pun1’, and ‘Sy-2’), were used in this study. *C. annuum* lines ‘106’ and ‘Nikko’ were crossed to generate an F_1_ mapping population. Then, the *C. chinense* line ‘pun1’ was crossed with a ‘106’ × ‘Nikko’ F_1_ plant to obtain a mapping population (*n* = 118).

### Genome sequencing and data analysis

A short-read sequence library of ‘Takanotsume’ was prepared using the TruSeq DNA PCR-Free Sample Preparation Kit (Illumina) and sequenced on the NextSeq500 instrument (Illumina) in paired-end 151 bp mode. After removing low-quality bases (quality value < 10) with PRINSEQ^21^ and adaptor sequences (AGATCGGAAGAGC) with fastx_clipper in the FASTX-Toolkit (http://hannonlab.cshl.edu/fastx_toolkit), the genome size of ‘Takanotsume’ was estimated using Jellyfish (*k*-mer size = 17)^22^.

### Linked-read sequencing and assembly

Genomic DNA was extracted from the young leaves of ‘Takanotsume’, ‘3686’, and ‘Sy-2’ plants using Genomic Tip (Qiagen, Hilden, Germany), and high-molecular-weight DNA (fragment length > 40 kb) was selected with BluePippin (Sage Science, Beverly, MA, USA). Genomic DNA library was prepared using the Chromium Genome Library Kit v2 (10X Genomics, Pleasanton, CA, USA) and sequenced on the NovaSeq 6000 platform (Illumina, San Diego, CA, USA) to generate paired-end 150 bp reads. The obtained sequence reads were assembled with Supernova (10X Genomics).

### Long-read sequencing and assembly

The genomic DNA of ‘Takanotsume’ used for linked-read sequencing was also used for long-read sequencing. Briefly, the genomic DNA of ‘Takanotsume’ was sheared in a DNA Shearing Tube g-TUBE (Covaris, Woburn, MA, USA) by centrifugation at 1,600 × g. The sheared DNA was used for HiFi SMRTbell library preparation with the SMRTbell Express Template Prep Kit 2.0 (PacBio, Menlo Park, CA, USA). The resultant library was separated on BluePippin (Sage Science) to remove short DNA fragments (<20 kb), and sequenced with SMRT cell 8 M on Sequel II and Sequel IIe systems (PacBio). The obtained HiFi reads were assembled with Hifiasm^23^.

### Optical mapping

Genomic DNA was extracted from young ‘Takanotsume’ leaves using the Plant DNA Isolation Kit (Bionano Genomics, San Diego, CA, USA), in accordance with Bionano Prep Plant Tissue DNA Isolation Base Protocol. The isolated genomic DNA was treated with DLE-1 nickase, and labeled with a florescent dye supplied in the DLS DNA Labeling Kit (Bionano Genomics). The labeled DNA was run on the Saphyr Optical Genome Mapping Instrument (Bionano Genomics). The output reads were assembled and then merged with the HiFi assembly to generate hybrid scaffold sequences using Bionano Solve (Bionano Genomics).

### Genetic mapping and chromosome-level assembly

Genomic DNA was extracted from all F_2_ individuals (*n* = 118) and their parental lines using the DNeasy Plant Mini Kit (Qiagen). The obtained DNA samples were digested with *Pst*I and *Msp*I to construct a double digest restriction-site associated DNA sequencing (ddRAD-Seq) library^24^, which was sequenced on HiSeq 4000 (Illumina) in paired-end mode. The obtained sequence reads were subjected to quality control (as described above), and mapped onto the hybrid scaffold sequences with Bowtie 2^25^. High-confidence biallelic single nucleotide polymorphisms (SNPs) were identified using the mpileup option of SAMtools^26^, and filtered using VCFtools^27^ using the following criteria: read depth ≥ 5; SNP quality = 10; proportion of missing data < 50%. The identified SNPs were subjected to linkage analysis using Lep-Map3^28^. Contig sequences were anchored to the genetic map, and pseudomolecule sequences were established with ALLMAPS^29^. Using D-genies^30^, the genome structure of ‘Takanotsume’ was compared with those of four *C. annuum* lines (‘CM334’ [GenBank accession no.: AYRZ00000000], ‘Zunla-1’ [ASJU00000000], ‘UCD-10X-F1’ [NPHV00000000], and ‘CA59’ [JAJQWV000000000]), one *C. baccatum* line (‘PBC81’ [MLFT00000000]), and one *C. chinense* line (‘PI159236’ [MCIT00000000]).

### Gene and repeat prediction

Gene prediction was performed with BRAKER2^31^, based on the peptide sequences of the predicted genes of *C. annuum* line ‘CM334’ and RNA-Seq reads obtained from the Sequence Read Archive (SRA) database of the National Center of Biotechnology Information (NCBI) (accession nos.: SRR17837286–SRR17837292 and SRR17837303–SRR17837315). Simultaneously, gene sequences reported in the genome assemblies of *C. annuum* lines ‘CM334’ and ‘Zunla-1’ were mapped onto the ‘Takanotsume’ genome assembly with Liftoff^32^. Genome positions of the predicted and mapped genes were compared with BEDtools^33^.

Repetitive sequences in the genome assembly of ‘Takanotsume’ were identified with RepeatMasker (https://www.repeatmasker.org) using repeat sequences registered in Repbase and a *de novo* repeat library built with RepeatModeler (https://www.repeatmasker.org).

### Genetic diversity analysis

Whole-genome shotgun libraries of 13 *Capsicum* lines were prepared with the TruSeq DNA PCR-Free Sample Prep Kit (Illumina), in accordance with the manufacturer’s protocol. The resultant libraries were sequenced either on HiSeq 2500 (Illumina) to generate paired-end 250 bp reads or on NextSeq500 (Illumina) and NovaSeq 6000 (Illumina) platforms to generate paired-end 151 bp reads. The reads were subjected to quality control (as described above) and mapped onto the pseudomolecule sequences of ‘Takanotsume’ with Bowtie2^25^. Sequence variants were detected using the mpileup and call commands of BCFtools^26^, and high-confidence biallelic SNPs were identified with VCFtools^27^ using the following parameters: minimum read depth ≥ 8 (--minDP 8); minimum variant quality = 20 (--minQ 20); maximum missing data < 0.5 (--max-missing 0.5); and minor allele frequency ≥ 0.05 (--maf 0.05). Effects of SNPs on gene function were estimated with SnpEff^34^. The population structure of the 13 *Capsicum* lines and ‘Takanotsume’ were evaluated with maximum likelihood estimation of individual ancestries with ADMIXTURE^35^ and principal component analysis with TASSEL^36^.

## Results

### Assembly of Capsicum genomes

Genome size estimation with 74.7 Gb short-read data indicated that ‘Takanotsume’ has a homozygous genome, with an estimated haploid genome size of 3,168.4 Mb (Figure 1).

**Figure 1.**
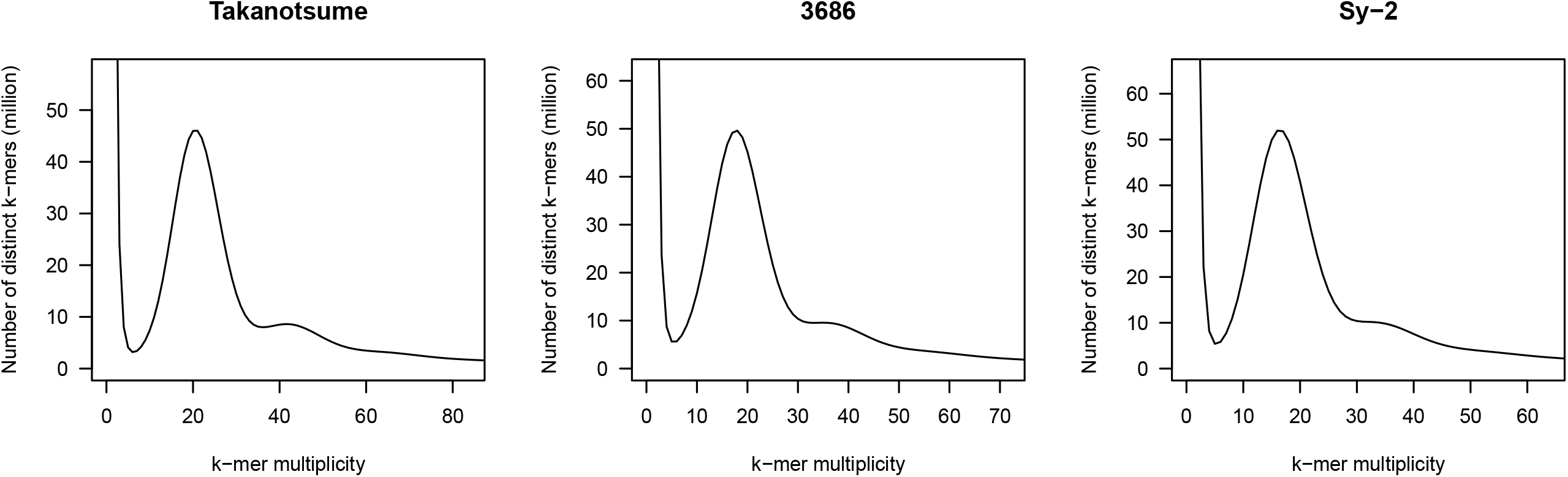
Estimated genome sizes of *Capsicum annuum* landrace ‘Takanotsume’ and *C. chinense* lines ‘3686’ and ‘Sy-2’, based on *k*-mer analysis (*k*□=□17), with the given multiplicity values.

The linked reads of ‘Takanotsume’ (497.2 Gb) were assembled into contigs, resulting in a total of 30,425 sequences (total length = 3,072.8 Mb; contig N50 length = 9.4 Mb) (Table 1). The linked-read sequencing-based genome assembly of ‘Takanotsume’ was designated as CAN_r0.1. The complete BUSCO score was 96.9%; however, the contigs were fragmented and exhibited short sequence contiguity.

**Table 1.** Statistics of the genome assemblies of three *Capsicum* lines belonging to two species.

|  | *C. annuum* |  |  |  | *C. chinense* |  |
| --- | --- | --- | --- | --- | --- | --- |
|  | ‘Takanotsume’ |  |  |  | ‘3686’ | ‘Sy-2’ |
| Sequencing technology | Linked-read | HiFi | HiFi + optical mapping | HiFi + optical mapping + genetic mapping | Linked read | Linked read |
| Total contig size (bp) | 3,072,766,948 | 3,094,642,642 | 3,074,206,442 | 3,058,489,554 | 3,019,584,847 | 3,000,496,031 |
| No. of contigs | 30,425 | 610 | 23 | 12 | 31,863 | 30,812 |
| Contig N50 length (bp) | 9,406,601 | 99,049,140 | 253,267,520 | 262,665,162 | 8,977,441 | 12,719,824 |
| Longest contig size (bp) | 61,314,963 | 177,899,727 | 337,042,926 | 337,042,926 | 45,657,838 | 74,881,574 |
| Gap (bp) | 60,035,580 | 0 | 8,701,469 | 8,181,982 | 60,254,330 | 60,914,730 |

To improve the ‘Takanotsume’ genome assembly, we employed the HiFi long-read sequencing technology. Five SMRT cells were used, generating 3,127,118 HiFi reads (total length = 66.0 Gb; N50 length = 21.4 kb; genome coverage = 20.8X). The reads were assembled into 610 primary contigs (total length = 3,094.6 Mb; N50 = 99.0 Mb) (Table 1). The complete BUSCO score was 97.4%, of which 95.6% were single-copy BUSCOs. The long-read-based genome assembly of ‘Takanotsume’ was designated as CAN_r1.0.

To extend the sequence contiguity, optical mapping was performed. Data amounting to 1,201.0 Gb (read length ≥ 150 kb) were generated, a subset (600 Gb) of which were employed for further analysis. Of the 600 Gb data, 563.8 Gb data (number of reads = 1,355,894; N50 length = 407.5 kb) were used for *de novo* assembly, generating 40 molecule maps (total length = 3,078.9 Mb; N50 = 247.1 Mb). In the subsequent hybrid scaffolding process, 2 and 16 conflicts in the 40 molecule maps and CAN_r1.0, respectively, were resolved. Then, a hybrid scaffold comprising 23 sequences (total length = 3,074.2 Mb; N50 = 253.3 Mb) was obtained (Table 1), which was designated as CAN_r1.1.

To anchor the CAN_r1.1 scaffold sequences to the chromosome, genetic mapping was performed. The DNA of the mapping population and their parental lines was subjected to ddRAD-Seq analysis, which generated 1.0 M reads per sample. After quality filtering, high-quality reads were mapped onto the CAN_r1.1 assembly, with a mapping rate of 92.5%. This resulted in the detection of 1,836 high-confidence SNPs. A linkage analysis of these SNPs resulted in a genetic map, with a total of 12 linkage groups and 1,736 SNPs, and a total genetic distance of 748.8 cM (Table 2). Eighteen CAN_r1.1 scaffolds were anchored to the genetic map. Nine of these scaffolds were anchored to nine chromosomes (ch01, ch02, ch03, ch04, ch05, ch06, ch08, ch09, and ch10; one per chromosome), while the remaining nine scaffolds were anchored to ch11 (two scaffolds), ch12 (three scaffolds), and ch07 (four scaffolds). Multiple scaffolds anchored to a particular chromosome were concatenated with 100 Ns. In total, 12 pseudomolecules spanning 3,058.5 Mb were established (Table 1). This final assembly was designated as CAN_r1.2.pmol.

**Table 2.** Statistics of ‘Takanotsume’ pseudomolecule sequences.

| Chromosome | No. of<br>SNPs | Genetic distance<br>(cM) | No. of<br>contigs | Physical distance<br>(bp) |
| --- | --- | --- | --- | --- |
| 1 | 168 | 102.9 | 1 | 337,042,926 |
| 2 | 147 | 63.8 | 1 | 177,533,547 |
| 3 | 245 | 83.3 | 1 | 292,038,349 |
| 4 | 118 | 71.0 | 1 | 253,125,799 |
| 5 | 81 | 61.4 | 1 | 251,194,792 |
| 6 | 163 | 52.5 | 1 | 253,267,520 |
| 7 | 123 | 58.0 | 4 | 267,785,325 |
| 8 | 149 | 47.1 | 1 | 175,912,755 |
| 9 | 99 | 31.6 | 1 | 266,007,691 |
| 10 | 138 | 52.0 | 1 | 244,603,042 |
| 11 | 162 | 63.2 | 2 | 277,312,646 |
| 12 | 143 | 62.0 | 3 | 262,665,162 |
| **Total** | **1,736** | **748.8** | **18** | **3,058,489,554** |

### Gene and repeat prediction

A total of 102,153 protein-coding genes were predicted in the CAN_r1.2.pmol assembly. Genes from the previously established genome assemblies of ‘CM334’ and ‘Zunla-1’ (30,242 and 35,336, respectively) were aligned against the CAN_r1.2.pmol to compare the genomic positions of predicted genes. Of the total of 29,899 ‘CM334’ and 34,482 ‘Zunla-1’ genes mapped onto the CAN_r1.2.pmol, 24,724 ‘CM334’ and 31,206 ‘Zunla-1’ genes coincided with the genomic positions of 34,324 of the total 102,153 predicted genes. These 34,324 genes and the remaining 67,829 genes were defined as high- and low-confidence genes, respectively. The complete BUSCO score of the high-confidence genes was 95.0%.

Repetitive sequences occupied a total physical distance of 2,549.4 Mb (83.4%) in the CAN_r1.2.pmol genome assembly (3,058.5 Mb). Nine major types of repeats were identified in varying proportions (Table 3). The dominant repeat types in the chromosome sequences were long-terminal repeats (LTRs; 63.1%, 1,928.4 Mb) including *Gypsy*- (54.0%, 1,651.9 Mb) and *Copia*-type retroelements (5.9%, 178.9 Mb). Repeat sequences unavailable in public databases totaled 273.0 Mb.

**Table 3.**
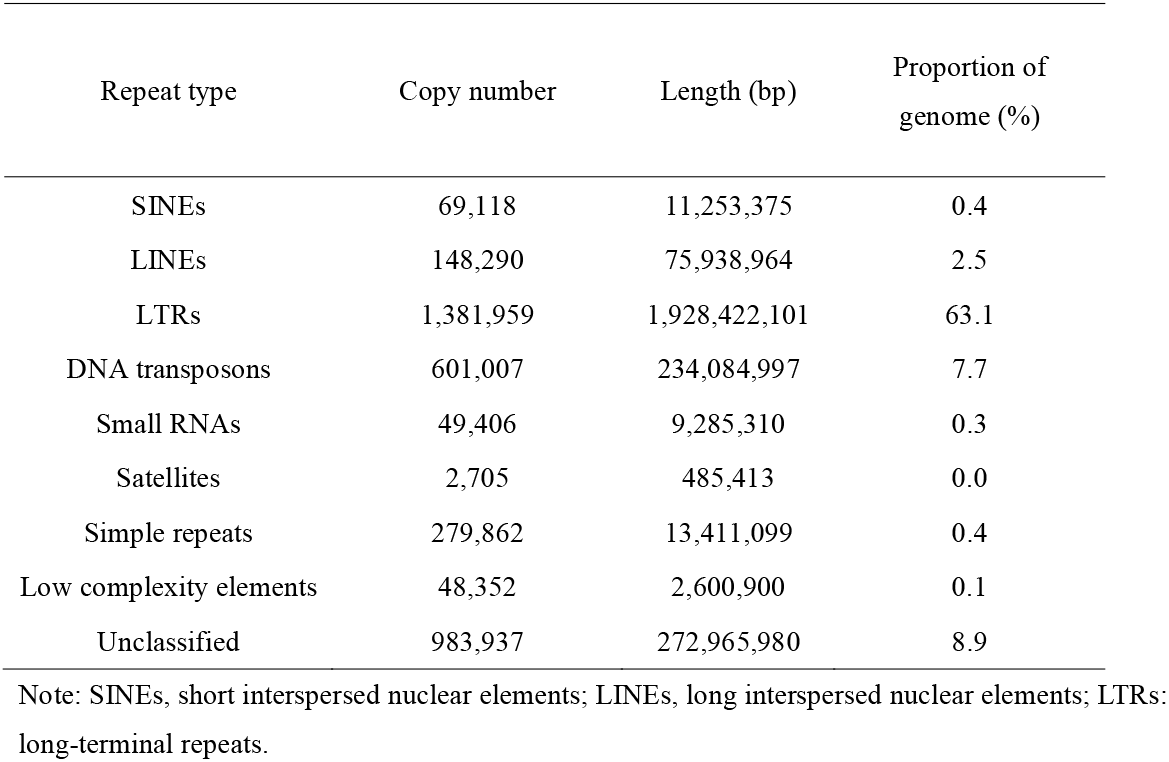
Repetitive sequences in the ‘Takanotsume’ genome.

### Sequence and structural variations within the genus Capsicum

First, genome structure variants between *C. annuum* and *C. chinense* were investigated. Genome sequences of two *C. chinense* lines, ‘3686’ and ‘Sy-2’, were constructed with the linked-read technology. The genome of ‘3686’ was 3,211.8 Mb in size, according to the *k*-mer frequency analysis (Figure 1), and the resultant assembly was 3,019.6 Mb in size, with 31,863 sequences and a contig N50 length of 9.0 Mb (Table 1). On the other hand, the genome size of ‘Sy-2’ was estimated as 3,303.2 Mb (Figure 1), and the assembly size was 3,000.5 Mb, including 30,812 sequences with a contig N50 length of 12.0 Mb (Table 1). The complete BUSCO scores of ‘3686’ and ‘Sy-2’ genomes were 96.0% and 96.6%, respectively. Finally, alignment analysis revealed that the ‘3686’, ‘Sy-2’, and CAN_r0.1 sequences covered 85.6%, 85.2%, and 96.3% of the CAN_r1.2.pmol reference sequence.

Next, sequence variants were detected six *C. annuum*, two *C. baccatum*, and five *C. chinense* lines. On average, 84.5 Gb short-read data were obtained from the 13 lines, and mapped onto CAN_r1.2.pmol, with mapping rates of 96.4% for *C. annuum*, 80.2% for *C. baccatum*, and 87.3% for *C. chinense*. Totals of 5.2, 32.9, and 43.8 million high-confidence SNPs were found in *C. annuum, C. baccatum*, and *C. chinense*, respectively. In the *C. annuum* lines, the SNP distribution pattern was biased (Figure 2), with a high density on ch09, ch10, and ch11 of ‘106’, ‘110’, ‘California Wonder’, and ‘Sweet Banana’. According to SnpEff results, the most prominent SNP type was modifier impact (98.5%) in intergenic regions and introns, followed by moderate impact (0.9%; leading to missense mutations), low impact (0.5%; synonymous mutations), and high impact (0.1%; nonsense mutations and mutations at splice junctions) (Table 4). The admixture analysis indicated the 13 lines in addition to ‘Takanotsume’ were grouped into three clusters (K) (Figure 3A): a) seven *C. annuum* lines, b) one *C. battatum* line, and c) the remaining five *C. chinense* lines in addition to one ‘Aji Roji’ (*C. baccatum*) (Figure 3B). When the number of K was increased to four, the seven *C. annuum* lines were separated into two groups (Figure 3B): 1) ‘Takanotsume’, ‘Nikko’, and ‘Murasaki’ and 2) ‘106’, ‘110’, ‘California Wonder’, and ‘Sweet Banana’. PCA analysis well supported the admixture result (Figure 3C).

**Table 4.**
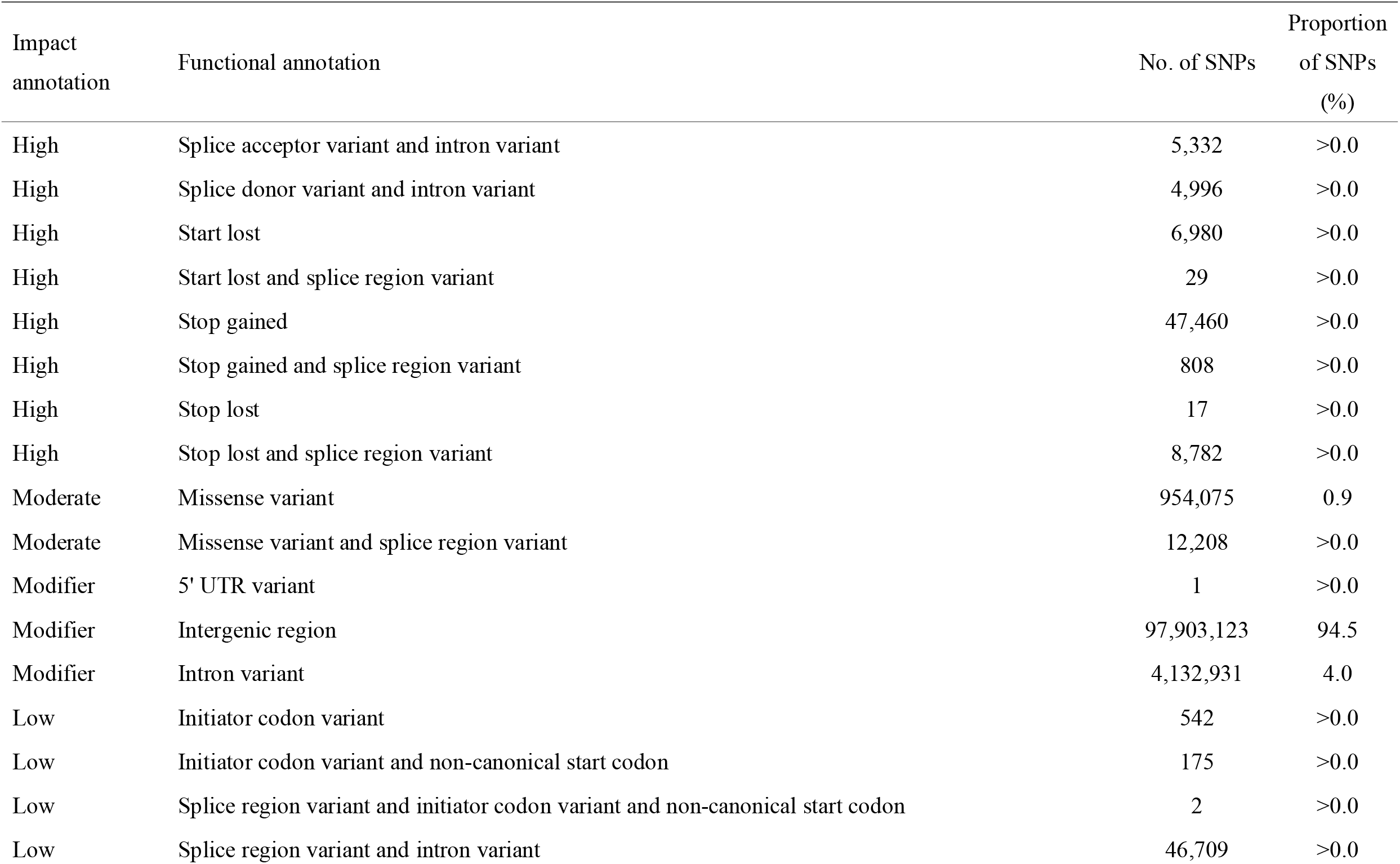
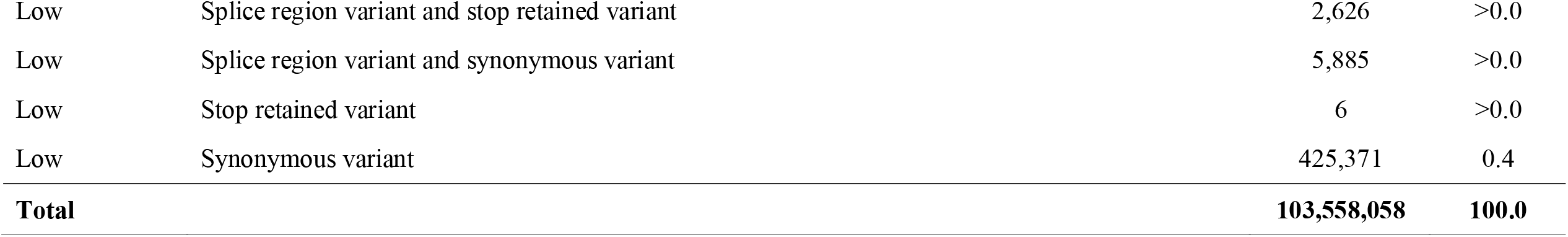
Annotation of variants detected among 13 lines of three *Capsicum* species.

| Impact annotation | Functional annotation | No. of SNPs | Proportion of SNPs (%) |
| --- | --- | --- | --- |
| High | Splice acceptor variant and intron variant | 5,332 | >0.0 |
| High | Splice donor variant and intron variant | 4,996 | >0.0 |
| High | Start lost | 6,980 | >0.0 |
| High | Start lost and splice region variant | 29 | >0.0 |
| High | Stop gained | 47,460 | >0.0 |
| High | Stop gained and splice region variant | 808 | >0.0 |
| High | Stop lost | 17 | >0.0 |
| High | Stop lost and splice region variant | 8,782 | >0.0 |
| Moderate | Missense variant | 954,075 | 0.9 |
| Moderate | Missense variant and splice region variant | 12,208 | >0.0 |
| Modifier | 5' UTR variant | 1 | >0.0 |
| Modifier | Intergenic region | 97,903,123 | 94.5 |
| Modifier | Intron variant | 4,132,931 | 4.0 |
| Low | Initiator codon variant | 542 | >0.0 |
| Low | Initiator codon variant and non-canonical start codon | 175 | >0.0 |
| Low | Splice region variant and initiator codon variant and non-canonical start codon | 2 | >0.0 |
| Low | Splice region variant and intron variant | 46,709 | >0.0 |
| Low | Splice region variant and stop retained variant | 2,626 | >0.0 |
| Low | Splice region variant and synonymous variant | 5,885 | >0.0 |
| Low | Stop retained variant | 6 | >0.0 |
| Low | Synonymous variant | 425,371 | 0.4 |
| **Total** |  | **103,558,058** | **100.0** |

**Figure 2.**
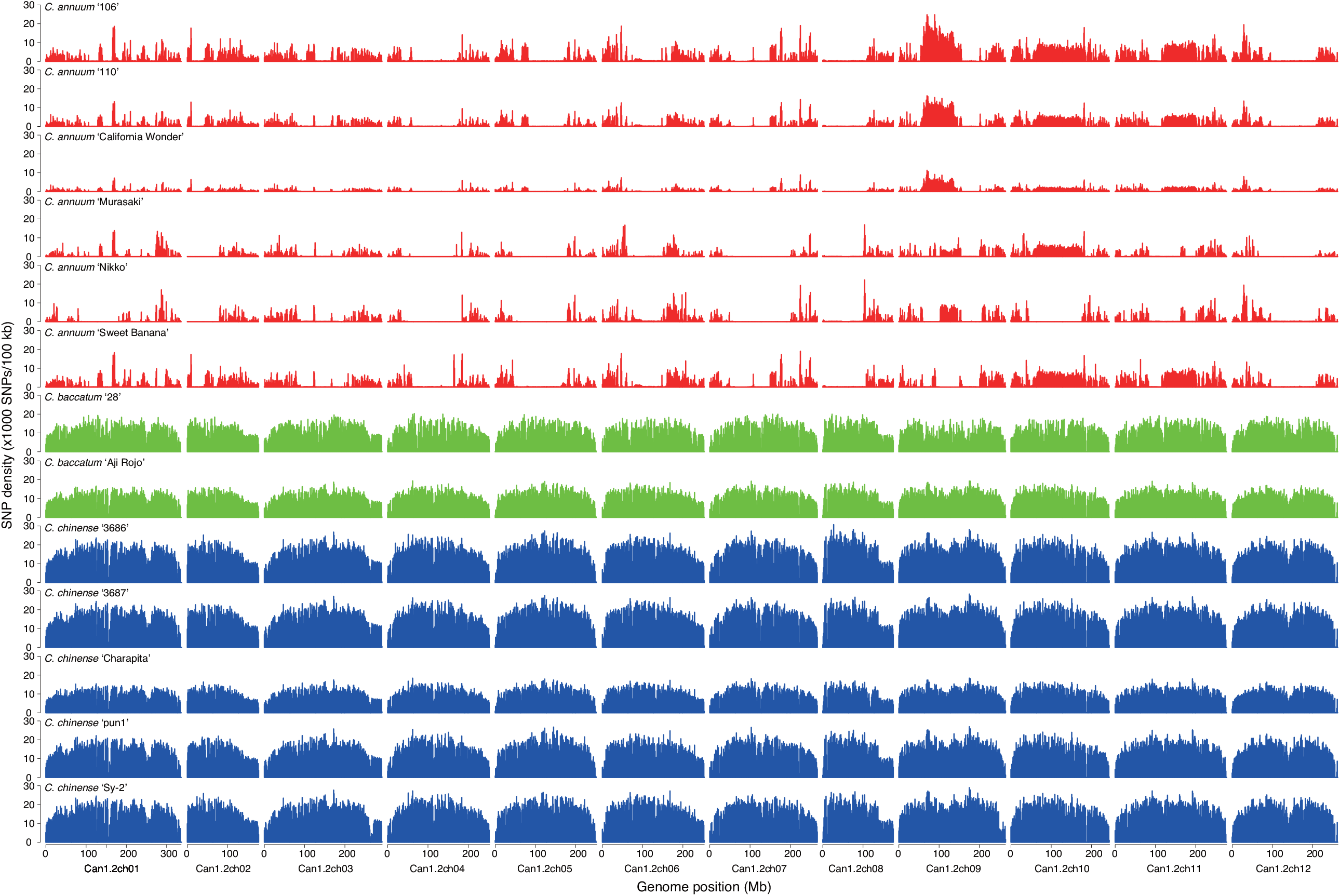
SNP density across the 13 *Capsicum* lines, based on the comparison with the ‘Takanotsume’ genome. *C. annuum, C. baccatum*, and *C. chinense* lines are shown in red, green, and blue, respectively.

**Figure 3.**
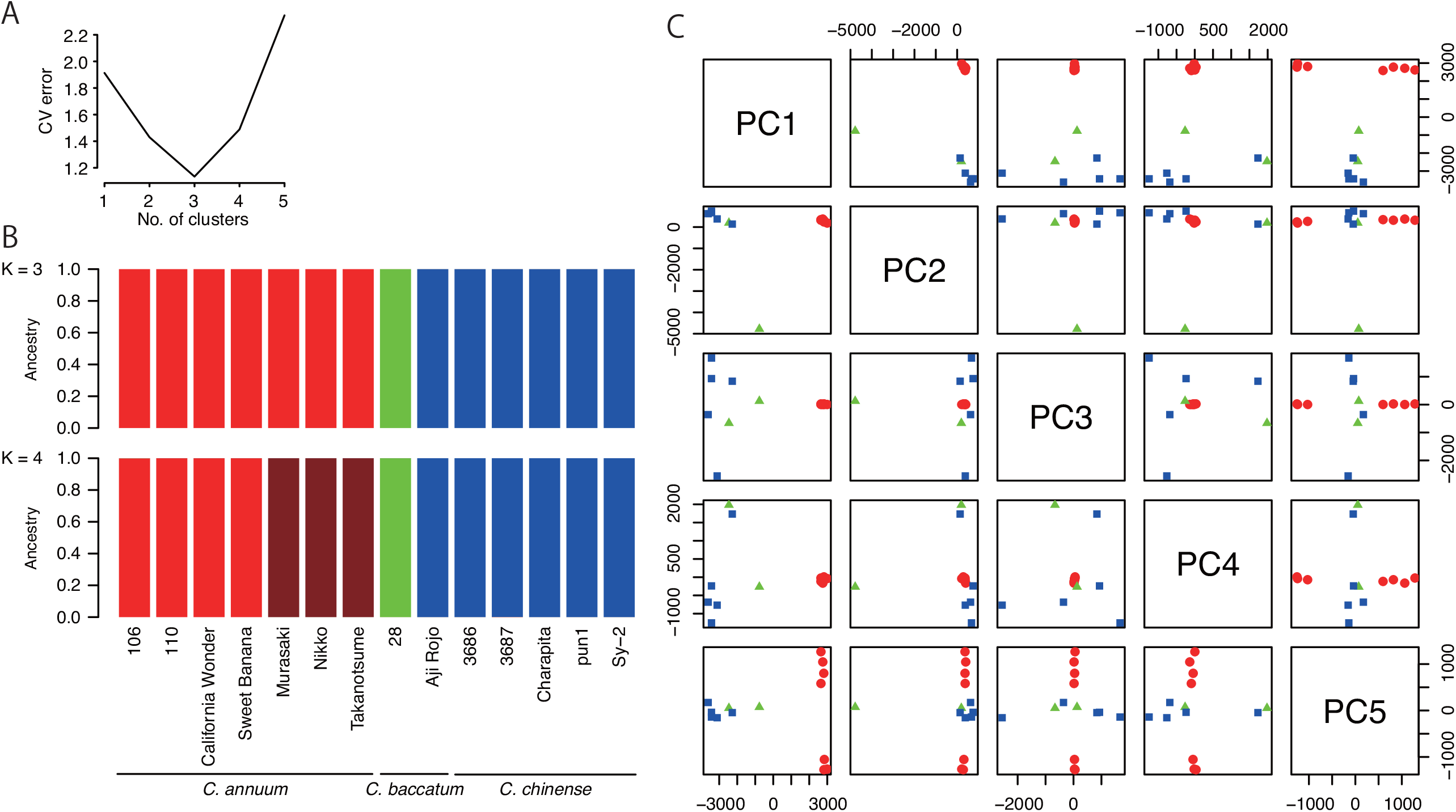
Genetic structure of the 14 *Capsicum* lines. **A**. Cross validation (CV) error plot for admixture analysis of K ranging from 1 to 5. **B**. Population structure of the 14 Capsicum lines. Each color represents a distinct group. **C**. Principal component analysis for the 14 Capsicum lines. *C. annuum, C. baccatum*, and *C. chinense* dots are shown in red, green, and blue, respectively.

Comparative genomics revealed that the genome structures of ‘Takanotsume’ and ‘CA59’ were well conserved; however, the chromosomes of five Capsicum lines (‘CM334’, ‘Zunla-1’, ‘UCD-10X-F1’, ‘PBC81’, and ‘PI159236’) were disrupted at the middle (Figure 4). Moreover, five potential translocations were detected in the ‘Takanotsume’ genome, including one on ch01 (compared with the ch08 of ‘PBC81’ and ‘PI159236’), two on ch03 (one compared with the ch05 of ‘PBC81’ and another relative to the ch09 of ‘PBC81’), one on ch05 (compared with the ch03 of ‘PBC81’), and one on ch09 (compared with the ch03 of ‘PBC81’).

**Figure 4.**
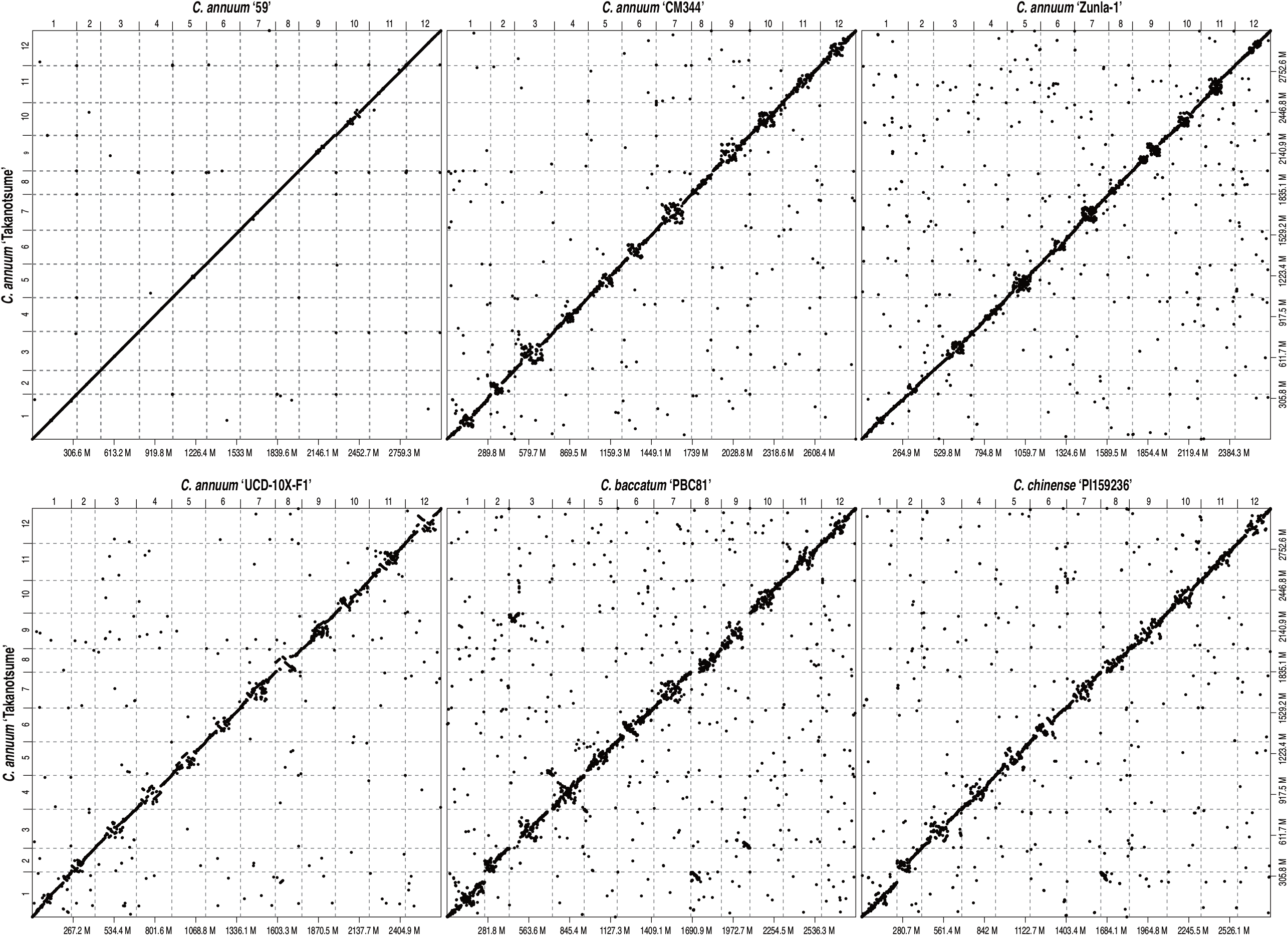
Comparative genomics of ‘Takanotsume’ and six divergent *Capsicum* lines belonging to three different species. Dots indicate structural similarities among the genomes of *Capsicum* species. Chromosome numbers are indicated above the x-axis and on the left-hand side of the y-axis, and genome sizes (Mb) are shown below the x-axis and on the right-hand side of the y-axis.

## Discussion

Here, we present the chromosome-scale genome assembly (CAN_r1.2.pmol) of a popular Japanese chili pepper *C. annuum* landrace, ‘Takanotsume’. The assembly spanned a total length of 3,058 Mb, which corresponded to 96.5% of the estimated genome size (Figure 1, Table 1). Sequence gaps (total length = 8.2 Mb) were observed at 171 locations on 12 chromosomes (Table 1). The contiguity of this chromosome-level assembly was much greater than that obtained using the 10XGenomics Chromium technology (Table 1). The genome coverage of the ‘Takanotsume’ assembly was comparable with that of ‘CA59’ and higher than those of ‘CM334’, ‘UCD-10X-F1’, ‘Zunla-1’, and the relatives ‘PBC81’ and ‘PI159236’. Moreover, sequence orders and orientations in the middle of the chromosomes were disrupted (Figure 4). This suggested that the genome structures varied within the *Capsicum* genus and/or there were misassembly points in the genomes of the five abovementioned lines, probably because of the short-read sequencing technologies employed. To validate this assumption, further karyotyping studies with fluorescence *in situ* hybridization are required. In addition, genic regions in the CAN_r1.2.pmol assembly were also well annotated (Table 3). A total of 34,324 high-confidence genes in CAN_r1.2.pmol were supported by those in ‘CM334’ and/or ‘Zunla-1’.

The population structure analyses indicated the three *Capsicum* species could be discriminated with the genetic variations of the genome (Figure 3) except for ‘Aji Rojo’, which is a *C. baccatum* line but grouped in the *C. chinense* cluster. In our previous studies^10,37^, 192 *Capsicum* lines mainly including four species, *C. annuum, C. baccatum, C. chinense*, and *C. frutescens*, were roughly classed into four groups representing the species; however, there were mismatch between the classifications based on morphological traits and those based on DNA sequence, probably due to misclassification of species based on morphological traits and/or genome introgression between different species^37^. These observations suggested that the concept of species might be reconsidered as discussed for a long time^38,39^, especially for crops including *Capsicum* because of the ease of the cross-compatibility between species. Indeed, in accordance with nuclear and plastid genotypes, ‘Takanotsume’ is suggested to be a derivative of the hybridization between *C. annuum* as a paternal parent and either *C. chinense* or *C. frutescens* as a maternal parent^37^. The ‘Takanotsume’ genome assembly from this study might contribute to clarify the mysterious pedigree.

‘Takanotsume’ exhibits attractive, agriculturally important phenotypes^18^. One of them is the restoration of hybrid breakdown in the progeny derived from crosses between *C. annuum* and either *C. baccatum* or *C. chinense*^3^. This phenomenon could be explained by the BDM model^4^, which was originally proposed >100 years ago; however, the molecular mechanisms still remain unclear. Owing to the high-quality genome assemblies and high coverage of the gene-rich regions, a map-based cloning strategy, together with gene editing and/or virus-induced gene silencing, would identify the genes capable of restoring hybrid breakdown in pepper. This would provide new insights into the molecular mechanisms responsible for the long-term unresolved BDM model. Another important characteristic of ‘Takanotsume’ is high ribonuclease activity in leaves^20^. This trait would be agronomically useful for the development of biopesticides to combat RNA viruses around the world. Identification of the genes responsible for the RNase activity in ‘Takanotsume’ would enable the regulation of enzyme activity and specificity.

In addition to nucleotide sequence polymorphisms, structural variations including copy-number variations (also known as presence-absence variations) and chromosomal rearrangements (such as translocations and inversions) can also explain the within-species phenotypic variation. Therefore, a single reference genome sequence of a species is insufficient for gaining insights into its genomics and genetics^40^. A genome sequence established by sequencing the genomes of multiple lines of a species is called the pan-genome^41^. Although a pan-genome study of *Capsicum* was recently conducted^17^, the corresponding genome sequence database was unavailable during the writing of this manuscript. In the future, pan-genome databases will be made publicly available, which will likely accelerate the pace of *Capsicum* genomics. The chromosome-level genome assembly of ‘Takanotsume’ constructed in this study is expected to contribute to the pan-genome study of *Capsicum*.

## Data availability

Raw sequence reads were deposited in the Sequence Read Archive (SRA) database of the DNA Data Bank of Japan (DDBJ) under the accession numbers DRA014624 and DRA014640–DRA014642. The assembled sequences are available at DDBJ (accession numbers AP026696–AP026707) and Plant GARDEN (https://plantgarden.jp).

## Acknowledgments

We thank Y. Kishida, C. Minami, K. Ozawa, H. Tsuruoka, and A. Watanabe (Kazusa DNA Research Institute) for technical assistance. This study was supported in part by JSPS KAKENHI (16H02535, 16H06279, 20H02981, 22H05172, and 22H05181) and the Kazusa DNA Research Institute Foundation.

